# Testing a candidate meiotic drive locus identified by pool sequencing

**DOI:** 10.1101/2023.05.04.539432

**Authors:** Daniel A. Barbash, Bozhou Jin, Kevin H-C Wei, Anne-Marie Dion-Côté

## Abstract

Meiotic drive biases the transmission of alleles in heterozygous individuals, such that Mendel’s law of equal segregation is violated. Most examples of meiotic drive have been discovered over the past century based on causing sex-ratio distortion or the biased transmission of easily scoreable genetic markers that were linked to drive alleles. More recently, several approaches have been developed that attempt to identify distortions of Mendelian segregation genome-wide. Here we test a candidate female meiotic drive locus in *Drosophila melanogaster*, identified previously as causing a ∼54:46 distortion ratio using sequencing of large pools of backcross progeny. We inserted fluorescent visible markers near the candidate locus and scored transmission in thousands of individual progeny. We observed a small but significant deviation from the Mendelian expectation, however it was in the opposite direction to that predicted based on the original experiments. We discuss several possible causes of the discrepancy between the two approaches, noting that subtle viability effects are particularly challenging to disentangle from potential small-effect meiotic drive loci. We conclude that pool sequencing approaches remain a powerful method to identify candidate meiotic drive loci, but that genotyping of individual progeny at early developmental stages may be required for robust confirmation.

## Introduction

Mendel’s first law, that heterozygous individuals transmit each of their two alleles at equal frequency, is well established. But rare cases of meiotic drivers that generate exceptions to this law have been found in many organisms, with some alleles biasing their own transmission almost completely. While meiotic drive has become a coverall term for a range of mechanisms causing transmission ratio distortion, we distinguish here two distinct processes that cause such distortion: “True” meiotic drive in females and gamete competition in males.

“True” meiotic drive can only occur in an animal or plant that has an asymmetric meiosis where one meiotic product becomes an egg or ovum while the other meiotic products are discarded as polar bodies (Novitski 1967; Pardo-Manuel de Villena and Sapienza 2001a; Chmátal *et al*. 2017). If an allele in a heterozygote can bias its transmission to the egg versus a polar body, then it will have a direct advantage and increase in frequency. Examples include knobs in maize, Robertsonian translocations in animals, structurally variant chromosomes in Drosophila, and centromeric alleles in Mimulus and mice (Rhoades 1942; Novitski 1967; Pardo-Manuel de Villena and Sapienza 2001b; Fishman and Saunders 2008; Iwata-Otsubo *et al*. 2017).

In contrast, male meiosis is typically symmetric, with all four meiotic products becoming gametes. There is thus no opportunity for alleles to bias their transmission during meiosis. However, competition can occur post-meiotically between gametes carrying different alleles. Such processes would best be described as segregation distortion but are also often termed as meiotic drive (Sandler and Novitski 1957; Lindholm *et al*. 2016). It has been most readily detected as causing sex-ratio distortion when driving alleles are located on sex chromosomes, and several autosomal distorters have also been characterized (Jaenike 2001; Herrmann and Bauer 2012; Larracuente and Presgraves 2012). Analogous systems also occur in fungi where they manifest as spore killers (Zanders and Johannesson 2021).

On the one hand, meiotic drivers bias their own transmission and will quickly go to fixation if left unchecked. On the other hand, if meiotic drive alleles have pleiotropic fitness costs, they will induce selective pressure for suppressors, potentially leading to recurrent cycles of new drive alleles arising followed by host evolution (Sandler and Novitski 1957; Crow 1991). Recurrent cycles of meiotic drive and suppression have been proposed to be a major contributor to karyotype evolution, speciation, sex chromosome turnover, and host-protein evolution (Buckler *et al*. 1999; Zwick *et al*. 1999; Henikoff *et al*. 2001; Pardo-Manuel de Villena and Sapienza 2001b; Meiklejohn and Tao 2010).

Most meiotic drivers have been discovered fortuitously, in crosses that happened to have linked markers present that showed very large deviations or by causing significant skewing of sex ratios. This leaves important questions open of how common meiotic drive is in natural populations and whether moderate strength drive alleles may be more prevalent than currently appreciated. Detection of small or moderate strength alleles can be particularly challenging to distinguish from viability effects, which are almost certainly more prevalent than meiotic drive.

New methods are needed to address these questions, particularly approaches that provide genome-scale detection. For male meiotic drive, sequencing large pools of motile sperm or pollen is a promising way to directly type gametes and look for unequal prevalence of alleles (Corbett-Detig *et al*. 2015, 2019). Sequencing large pools of progeny is another approach applicable to inferring either male or female meiotic drive, or transmission-ratio distortion in general (Bélanger *et al*. 2016; Seymour *et al*. 2019; Ren *et al*. 2021). We previously developed one such method, where F1 progeny are made between two fully sequenced inbred strains of Drosophila, and then backcrossed to either one of the parental strains (Wei *et al*. 2017).

Assuming Mendelian segregation, any SNPs that distinguish the two strains will either be heterozygous or homozygous in first-generation backcross (BC1) progeny for SNPs derived from the parent used to backcross. If many progeny are pooled together and sequenced, then one expects to detect a 75:25 ratio of parental alleles, with deviations indicating potential meiotic drive.

We performed several sets of crosses and sequenced pools of thousands of BC1 progeny to screen for female meiotic drive loci in crosses between several *D. melanogaster* strains (Wei *et al*. 2017). In crosses with strains DGRP-129 and DGRP-882, we identified a locus mapping broadly to the chromosome 3 centromere region that confers an approximately 54:46 ratio in favor of the DGRP-882 allele. This result was replicated in both embryonic and adult progeny, suggesting that it is unlikely to be due to post-embryonic viability effects. The allele skew was only observed in the progeny of heterozygous females, not males, providing further evidence against viability effects and also strongly suggesting that it reflects female-specific meiotic drive.

As noted in that study, pooling has its caveats. Most importantly, any pooling strategy depends on the assumption that all individuals contribute equally to the DNA pool that is sequenced. This assumption would be violated if certain genotypes have more cells or have higher levels of polytenized nuclei; if so they will disproportionately contribute DNA to the pool in a way that mimics the effects of meiotic drive.

The most direct way to distinguish true inference of meiotic drive is to genotype individual progeny, which provides an independent test not relying on pooling. Here we do so for progeny of the same cross between DGRP-129 and DGRP-882.

## Results

We wished to insert easily scorable fluorescent markers that express both early in development and in adults, near the putative meiotic drive region of the chromosome 3 centromeres of both candidate strains. A *dfd-GMR* construct provides expression in embryos and in the adult proboscis via the *dfd* promoter and in adult eyes via the *GMR* promoter (Le *et al*. 2006). We replaced the GMR promoter with a 3XP3 promoter to obtain higher expression in the eyes and made versions expressing either Gfp or mCherry.

We introduced *dfd-3XP3-Gfp* into DGRP-882 and *dfd-3XP3-mCherry* into DGRP-129 at a pericentromeric site on chromosome 3 using CRISPR/Cas9-mediated recombination. Both transgenes expressed well (Fig 1). Signal in the eye driven by 3XP3 was difficult to detect at low magnification due to wild type eye pigmentation. We therefore relied on expression in the proboscis to score adults. We additionally observed high expression of mCherry in the abdomen, which is driven by the 3XP3 promoter (Champer *et al*. 2017). A similar signal cannot be detected with Gfp due to the high background in abdominal tissues in non-Gfp flies (Fig 1C).

**Figure 1.**
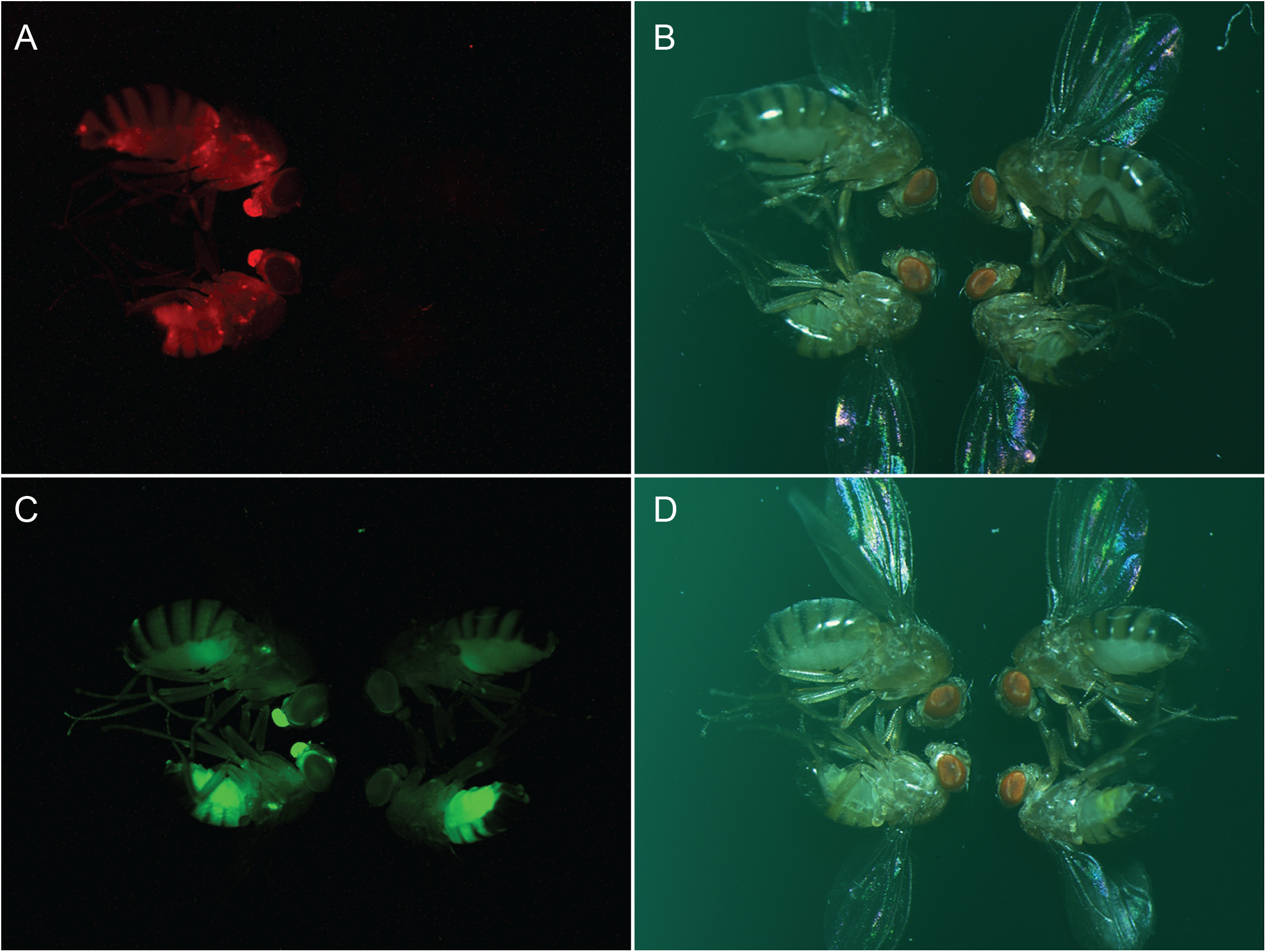
Strong expression of Gfp and mCherry in DGRP strains heterozygous for transgenes in the 3R pericentromere. Each panel shows a female (top) and male (bottom) carrying the transgene on the left side, and the same but not carrying the transgene on the right side. A) Image of mCherry signal and B) image under bright light of flies from a cross with DGRP-129 *mCherry*. C) Image of Gfp signal and D) image under bright light of flies from a cross with DGRP-882 *Gfp*. Wild-type females (Zambia line 379) were crossed separately to either DGRP-129 *mCherry* (A, B) or DGRP-882 *Gfp* (C, D) males at 25° C. F1 progeny were collected within 24 hours of eclosion, aged for 4 days, and then frozen at -80° C. Flies were thawed at room temperature and imaged using an Olympus SZX7 Stereo Microscopes connected to an Olympus America S97809 Camera.

We were able to establish and maintain lines that are viable and fertile as homozygotes for each of the two resulting strains. It was important though to determine whether either marker has any viability effect, as that could impair its use in accurately assessing a candidate meiotic driver of modest effect.

We therefore tested for potential dominant effects of each marker within its strain background, using a *white* mutant background for DGRP-882 in order to more easily score 3XP3-Gfp. We found that mCherry had no effect on viability in DGRP-129 but that Gfp caused a small but significant reduction in DGRP-882 (Table 1).

**Table 1.** Testing markers for potential viability effects

| Marker | No. with marker | No. without marker | No. with marker/total | p-value |
| --- | --- | --- | --- | --- |
| Gfp | 3463 | 3678 | 0.485 | 0.01132 |
| mCherry | 1693 | 1691 | 0.500 | 0.9863 |

We next tested for potential meiotic drive in heterozygous DGRP-129-mCherry^+^ / DGRP-882-Gfp^+^ F1 females (Table 2). We performed two sets of crosses, backcrossing the F1 females to DGRP-129 and DGRP-882 males, respectively. By scoring 6,027 and 4,672 progeny in the two sets, respectively, we were able potentially to detect small deviations from Mendelian expectations. While we found no significant differences in progeny ratios between the two sets of crosses, both sets gave small but significant deficits in the expected number of Gfp^+^ progeny, with ratios of 48.6% and 47.4%, respectively. Note that this deviation is opposite in direction to our hypothesis that DGRP-882 contains a mild female meiotic driver when heterozygous to DGRP-129.

**Table 2.** Crosses to test for meiotic drive in DGRP-129/DGRP-882 heterozygotes

| Sire | No. Gfp+ | No. mCherry+ | Ratio Gfp+/Total | *p*-value |
| --- | --- | --- | --- | --- |
| DGRP-129 | 2931 | 3096 | 0.486 | 0.0346 |
| DGRP-882 | 2216 | 2456 | 0.474 | 0.00047 |

This small deficit of Gfp^+^ progeny in our experimental crosses is very similar to the 48.5% reduction in Gfp^+^ progeny in our control crosses (Table 1). Assuming that the expression of Gfp does not have variable effects on viability in different genetic backgrounds, the most straightforward conclusion is that the small but significant deviations we observed in the experimental crosses are due to viability effects, not meiotic drive.

## Discussion

What then accounts for the different results here compared to Wei et al. (2017)? Even subtle effects could have a strong impact, given the relatively low magnitude of the putative meiotic drive locus reported in Wei et al. (2017). We consider four possibilities.

1. Environmental effects. The experiments here were performed several years after those in Wei et al. (2017), with possible differences in Drosophila media and building environment occurring.
2. Genetic drift. We confirmed the identity of the DGRP stocks during this study using diagnostic RFLPs, but that does not preclude the possibility that modifier alleles could have evolved over the several years in which these stocks were maintained between the two studies.
3. Marker variability. The fluorescent markers used here to genotype the individual flies had strong signals that we found straightforward to score. Given that they are inserted in pericentric heterochromatin, it is nevertheless possible that their expression might be variable or even suppressed in a small proportion of flies. It is also possible that expression is variable in some genotypes but not others. Variable expression in the experimental progeny that are heterozygous, relative to expression within the pure-strain homozygous backgrounds, would lead to mis-inferences.
4. Background-dependent viability and developmental effects. Viability and developmental effects that vary across genetic backgrounds could cause discrepancies similar to those described above for marker variability. In this study, we found that the *dfd-3XP3-Gfp* marker causes a slight reduction in viability in a pure DGRP-882 background (Table 1). Because we saw nearly the same deficit of Gfp^+^ progeny in the experimental crosses (Table 2), we concluded that there is no meiotic drive effect. This conclusion, however, assumes that any viability effect of the marker is constant regardless of background.

A broader range of potential developmental effects needs to be considered with the pooling approach used in Wei et al. (2017). Differences between genotypes in their number of cells and/or degree of polytenization could cause skews in allele frequencies that resemble meiotic drive effects. Wei et al. (2017) found that the allele skew only occurred in crosses where female parents were heterozygous but not in the reciprocal cross where male parents were heterozygous. This result was an important piece of evidence arguing for female meiotic drive rather than developmental effects. But as Wei et al. (2017) noted, it assumes that any developmental effects are constant across maternal and paternal genotypes. If instead developmental effects interact with or are caused by, for example, maternal effects, they could lead to mis-inference of meiotic drive.

Any potential effects that vary among genotypes are challenging to control for. We suggest that the least biased approach would be to genotype individual embryos, as viability effects are best minimized by genotyping as early in development as possible. Developing high-throughput, robust and inexpensive methods to genotype large numbers of embryos (at the earliest possible developmental stage) may provide the means to reliably screen for and test moderate-strength meiotic drive loci.

Nevertheless, we suggest that pooled sequencing methods remain a potentially powerful approach to identify loci causing meiotic drive, and more generally, transmission ratio distortion, due to their ability to screen the entire genome. It will though be important to confirm any candidates identified using a distinct method.

## Acknowledgements

We are grateful to Shuqing Ji for help performing crosses, and Dean Castillo and Sarah E. Lower for technical advice. This work was supported by National Institutes of Health (NIH) grant R01-GM07473 to D.A.B. A.M.D.C. was supported by postdoctoral fellowships from the FRQ-S (33616), NSERC (PDF-51651-2018) and the Lawski foundation as well as an NSERC Discovery grant (RGPIN-2019-0544).

## Materials and Methods

### Construction of Dfd-3XP3 marker plasmids

We amplified the 3xP3 promoter from pHD-DsRed-attP-w+ using oligos #1827 (contains a XhoI site; Table S1) and #1828 (contains a BglII site) and cloned it into pDf-GMR-nvYFP that had been digested with BamHI and XhoI to remove the GMR promoter (Le *et al*. 2006). pHD-DsRed-attP-w+ was a gift from Kate O’Connor-Giles (Addgene plasmid # 80898; http://n2t.net/addgene:80898; RRID:Addgene_80898). We then removed the Hsp70 promoter by digesting with PmeI and XhoI, forming blunt ends with Klenow enzyme treatment, gel purifying the fragment, phosphorylating with T4 polynucleotide kinase, self-ligating the product, and confirming the promoter region by sequencing. We next added an attB site by PCRing from pTA-attB (Groth *et al*. 2000) with oligos #502 and #503 (each of which contains a NotI site), digesting the product with NotI, and cloning into a unique NotI site that is 5’ to the *dfd* promoter, to create the plasmid pAttB-Dfd-3xP3-nvYFP. The plasmid was tested for expression by transforming into the strain *P{nos-phiC31\int*.*NLS}X, P{CaryP}attP2*.

We then replaced nvYFP with eGFP because GFP signal in the eye shows less quenching by wild-type eye pigmentation. We PCR-amplified eGFP from the plasmid pBS-KS-attB1-2-PT-SA-SD-0-EGFP-FlAsH-StrepII-TEV-3xFlag (Venken *et al*. 2011) using primers GFP-fwd and GFP-rev and PCR amplified attB-Dfd-3XP3 from pAttB-Dfd-3XP3-nvYfp with primers #1831 and #1832, then used Gibson assembly to make plasmid #758 = pAttB-Dfd-3xP3-GFP.

To replace nvYFP with mCherry we PCR amplified mCherry from pQC NLS mCherry IX (Addgene plasmid 37354) using primers #1833 and #1834 and PCR amplified attB-Dfd-3XP3 from pAttB-Dfd-3XP3-nvYfp with primers #1831 and #1832 and performed Gibson assembly to generate plasmid #795 = pAttB-Dfd-3xP3-mCherry.

### Construction of targeting plasmids

The plasmid #779 pHD-AttP-Dfd-3xP3-mCherry-3R was generated by Gibson assembly of the vector backbone from pHD-DsRed digested with EcoRI and XhoI and the following four PCR fragments: attP, amplified from p697 using oligos #1835 and #1836; promoter and reporter sequences amplified from pAttB-Dfd-3xP3-mCherry using oligos #1842 and #1843; left homology arm, amplified from DGRP-129 genomic DNA with oligos #1838 and #1839; right homology arm, amplified from DGRP-129 genomic DNA with oligos #1840 and #1841.

Plasmid #778 = pHD-AttP-Dfd-3xP3-eGFP-3R was similarly made except using pAttB-Dfd-3xP3-eGFP as the template for the reporter sequence and DGRP-882 genomic DNA as the template for the homology arms.

### Choosing target site and gRNA

We searched on Gbrowse for a region in the chromosome 3R pericentromere that is relatively free of repetitive DNA and contains transgenes with expressed visible markers, settling on 3R:4,150,200..4,150,600 (Flybase Release 6; Gramates *et al*. 2022). We chose a gRNA target site within this region at 4,150,207 - 4,150,226, with left and right homology arms at 4,148,824..4,150,224 and 4,150,224..4,151,624, respectively. The gRNA was cloned into pU6-BbsI-chiRNA (Gratz *et al*. 2013) using annealed oligos 1852 and 1853 to generate plasmid #773.

### Microinjection and strain construction

pHD-AttP-Dfd-3xP3-eGFP-3R plasmid along with Cas9-expressing plasmid pBS-Hsp70-Cas9 (Gratz *et al*. 2014) was injected into DGRP-882 by Genetivision. From 87 fertile G0 flies we obtained five candidate transformants from which one was confirmed to express *Gfp* and have the correct integration site. We refer to this strain as DGRP-882 *Gfp*.

pHD-AttP-Dfd-3xP3-mCherry-3R plasmid along with Cas9 protein was injected into DGRP-129 by Genetivision. From 131 fertile G0 flies we obtained three candidate transformants from which one was confirmed to express *mCherry* and have the correct integration site. We refer to this strain as DGRP-129 *mCherry*.

We confirmed the identity of the following strains immediately after performing the experiments in Table 2 and prior to those in Table 1: DGRP-129, DGRP-882, DGRP-129 *mCherry* and DGRP-882 *Gfp*. We used the RFLPs 2L5 and 3R2 that are diagnostic for these genotypes (MacKay et al. 2012).

### Generation of DGRP-882 *w* mutant

We designed two gRNAs to the *white* gene using the w1 and w2 sequences from (Ren *et al*. 2013), cloned into the plasmid pU6-BbsI-chiRNA. These two plasmids along with in vitro synthesized Cas9 mRNA were injected into DGRP-882 embryos by Rainbow Transgenic Flies. Two out of seventeen fertile G0 embryos produced white mutant progeny, which were brother-sister mated to produce a homozygous DGRP-882 *w* stock.

### Fly crosses

For the Gfp control crosses in Table 1, ∼15 DGRP-882 *w* virgin females were crossed to ∼15 DGRP-882 *w; Gfp* males. Two replicates of ∼15-20 DGRP-882 *w; Gfp/+* virgin daughters were then crossed to ∼15-20 DGRP-882 *w* males at 25° C. Vials were flipped every 1-2 days for a total of 6 flips. Similar crosses were done for the mCherry control, with DGRP-129 *mCherry* females crossed to DGRP-129 males, and then F1 daughters crossed to DGRP-129 males.

For the crosses in Table 2, two biological replicates were performed. Approximately 15-20 DGRP-882 *Gfp* females were crossed to ∼15-20 DGRP-129 *mCherry* males for each replicate. F1 virgin females were confirmed to express both Gfp and mCherry, and then crossed to males of either DGRP-129 or DGRP-882. Three sets were done with females aged 3-5 days and crosses set with 5 females and 5 males, with vials flipped every 1-2 days. As we found that many parents died by the third vial, we increased the number of parents to ∼10-14 of each sex for two more sets; the females for these sets were aged 8-11 days by happenstance. Only female progeny were scored, as was done in Wei et al. (2017).

DRGP-882 *w* females were crossed to DGRP-882 *Gfp* males. F1 DGRP-882 *w/Y; Gfp/+* males were mated to DRGP-882 *w* females. F2 DRGP-882 *w; Gfp/+* females were then mated to DRGP-882 *w* males and progeny scored.

DGRP-129 *mCherry* females were mated to DGRP-129 males. F1 DGRP-129 *mCherry/+* females were mated to DGRP-129 males and progeny scored. Significance of the ratio was tested using binom.test of the number of progeny carrying the marker relative to the total, given an expectation of 0.5 (R Core Team 2017).

DGRP-882 *Gfp* females were crossed to DGRP-129 *mCherry* males. F1 virgin females were then crossed to males of the genotype listed in column 1 (“Sire”). Significance of the ratio was tested using binom.test of the number of *Gfp*^*+*^ relative to the total, given an expectation of 0.5 (R Core Team 2017).

## Notes

### Competing Interest Statement

The authors have declared no competing interest.

