## Supplemental Table 1 for "Testing a candidate meiotic drive locus identified by pool sequencing"

Table S1. DNA primers used.

| Name | Sequence | Primer # |
|---|---|---|
| Not_attB_1 | aaacccgcggccgcatgcccgccgtgaccgtc | 502 |
| Not_attB_2 | aaacccgcggccgcgatgtaggtcacggtctcg | 503 |
| 3XP3-XhoI-R | TGCTGTCTCGAGGCCGATTGTTTAGCTTGTTCAGC | 1827 |
| 3xP3-BglII-F | GATCGAAGATCTGGATCTAATTCAATTAGAGACTAATTC | 1828 |
| pDfd_fwd | TAAGCGATCGCCTGATCATAATCAG | 1831 |
| pDfd_rev | GTTGACCGGTGGATCGTTTTCGAG | 1832 |
| mCherry_fwd | tcgaaaacgatccaccggtcaacATGGTGAGCAAGGGCGAG | 1833 |
| mCherry_rev | tatgatcaggcgatcgcttaTTACTTGTACAGCTCGTCCATGC | 1834 |
| 3R-LA_fwd | tgtcgcccttcgctgaagcaggtggCTTCATTATTCTCTACGTCTTG | 1838 |
| 3R-LA_rev | ccgctagcatgcaGATCTCAGAAACCATAATATCTATC | 1839 |
| AttP-short+plasmid_fwd | ggtttctgagatcTGCATGCTAGCGGCCGCG | 1835 |
| AttP-short+plasmid_rev | ctctagagcggccAGTCTCTAATTGAATTAGATCCCGTACGATAACTTCGTATA<br>GC | 1836 |
| Dfd-3xP3-nvYFP_fwd | tcaattagagactGGCCGCTCTAGAGAGGAC | 1842 |
| Dfd-3xP3-nvYFP_rev | catccgtcctgcgTAAGATACATTGATGAGTTTGGACAAAC | 1843 |
| 3R-RA_fwd | tcaatgtatcttaCGCAGGACGGATGGATGG | 1840 |
| 3R-RA_rev | cttgaactcgattgacggaagagccGTAAGCAACTTTAAAATACTCGTACAATTATTA<br>ACGC | 1841 |
| 3R sense | CTTCGATTATGGTTTCTGAGATCGC<br>CTTCGATTATGGTTTCTGAGATCGC | 1852 |
| 3R antisense | AAACGCGATCTCAGAAACCATAATC | 1853 |
| GFP_fwd | tcgaaaacgatccaccggtcaacATGGTGTCCAAGGGCGAGGAG | 1856 |
| GFP_rev | tatgatcaggcgatcgcttaCTACTTGTACAGCTCATCCATG | 1857 |
